# Nucleus accumbens melanin-concentrating hormone signaling promotes feeding in a sex-specific manner

**DOI:** 10.1101/2020.06.16.155747

**Authors:** Sarah J. Terrill, Keshav Subramanian, Rae Lan, Clarissa M. Liu, Alyssa M. Cortella, Emily E. Noble, Scott E. Kanoski

## Abstract

Melanin-concentrating hormone (MCH) is an orexigenic neuropeptide produced in the lateral hypothalamus and zona incerta that increases food intake. The neuronal pathways and behavioral mechanisms mediating the orexigenic effects of MCH are poorly understood, as is the extent to which MCH-mediated feeding outcomes are sex-dependent. Here we investigate the hypothesis that MCH-producing neurons act in the nucleus accumbens shell (ACBsh) to promote feeding behavior and motivation for palatable food in a sex-dependent manner. We utilized ACBsh MCH receptor (MCH1R)-directed pharmacology as well as a dual virus chemogenetic approach to selectively activate MCH neurons that project to the ACBsh. Results reveal that both ACBsh MCH1R activation and activating ACBsh-projecting MCH neurons increase consumption of standard chow and palatable sucrose in male rats without affecting motivated operant responding for sucrose, general activity levels, or anxiety-like behavior. In contrast, food intake was not affected in female rats by either ACBsh MCH1R activation or ACBsh-projecting MCH neuron activation. To determine a mechanism for this sexual dimorphism, we investigated whether the orexigenic effect of ACBsh MCH1R activation is reduced by endogenous estradiol signaling. In ovariectomized female rats on a cyclic regimen of either estradiol (EB) or oil vehicle, ACBsh MCH1R activation increased feeding only in oil-treated rats, suggesting that EB attenuates the ability of ACBsh MCH signaling to promote food intake. Collective results show that that MCH ACBsh signaling promotes feeding in an estrogen- and sex-dependent manner, thus identifying novel neurobiological mechanisms through which MCH and female sex hormones interact to influence food intake.

## 1. Introduction

MCH is a potent orexigenic neuropeptide produced by neurons in the lateral hypothalamic area (LHA) and zona incerta (ZI) (Barson et al., 2013; Qu et al., 1996; Shimada et al., 1998). Pharmacological activation of melanin-concentrating hormone receptor 1 receptor (MCH1R) both increases food intake and reduces energy expenditure, promoting overall elevated weight gain in rodents (Chambers et al., 1999; Kowalski et al., 2004; Lembo et al., 1999; Shearman et al., 2003). Mice lacking MCH (Shimada et al., 1998) or MCH1R (Marsh et al., 2002) are lean, whereas transgenic MCH overexpression produces hyperphagia and obesity (Ludwig et al., 2001). There is recent clinical interest in targeting the MCH system for obesity pharmacotherapy development (Abdel-Magid, 2015; Oost et al., 2015; Ploj et al., 2016), based in part on findings that chronic central MCH1R blockade reverses diet-induced obesity in mice partially through a reduction in caloric intake (Ito et al., 2010; Kowalski et al., 2006; Mashiko et al., 2005). While MCH1Rs are expressed throughout the neuraxis and MCH neurons have extensive projections (Chee et al., 2013; Diniz et al., 2019; Lembo et al., 1999), surprisingly little is known about the neurobiological and behavioral mechanisms through which MCH signaling promotes appetite and food intake.

The nucleus accumbens shell (ACBsh) is classically associated with drug addiction and reward-motivated behaviors, including the control of learned aspects of food reward (Alhadeff et al., 2012; Baldo and Kelley, 2007; Berridge et al., 2010; Castro et al., 2015; Dossat et al., 2011; Durst et al., 2019; Will et al., 2003). MCH1Rs are expressed in the ACBsh and pharmacological MCH delivery to the ACB increases chow intake during the dark cycle in male rats (Georgescu et al., 2005). However, the extent to which ACBsh MCH1R activation increases consumption of and motivation to work for palatable foods associated with obesity is poorly understood, as is whether ACBsh MCH1R signaling influences feeding in a sex-dependent manner. Moreover, under physiological conditions MCH1R-expressing neurons are not exclusively engaged by MCH, but also by various neurochemicals that are co-released by MCH-producing neurons (e.g., GABA, glutamate, CART) (Chee et al., 2015; Harthoorn et al., 2005; Mickelsen et al., 2017), and thus engaging MCH neurons may result in a different outcome than MCH1R specific pharmacological activation in the ACBsh. Whether activation of ACBsh-projecting MCH neurons affects feeding behavior has not been previously investigated.

Here, we focused on the ACBsh as a site of action for MCH effects on feeding in both male and female rats. We sought to determine whether intra-ACBsh MCH, at doses subthreshold for effect in the lateral ventricle, affects consumption of standard rat chow as well as palatable sucrose. Utilizing a dual virus chemogenetic approach, we further examined whether selective activation of ACBsh-projecting MCH neurons increases chow and/or sucrose intake. To elucidate behavioral mechanisms through which MCH signaling promotes appetite and food intake, we assessed the role of ACBsh MCH1R in effort-based responding for sucrose and evaluated the effects of selective activation of ACBsh-projecting MCH neurons on general activity levels in an open field procedure. Additional experiments investigate whether the female sex hormone, estradiol, modulates orexigenic effects of MCH1R activation in the ACBsh, as well as the extent that ACBsh MCH1R expression co-localizes with estrogen receptor differentially in male and female rats.

## 2. Methods

### 2.1. Subjects

Male Sprague Dawley rats (Envigo, Indianapolis, IN, USA) weighing 250–300 g and female Sprague Dawley rats (Envigo, Indianapolis, IN, USA) weighing 175-250 g were individually housed in wire-hanging cages in a climate controlled (22–24 °C) environment with a 12:12 h light/dark cycle. Except where noted, rats were given ad libitum access to water and standard rodent chow (LabDiet 5001, LabDiet, St. Louis, MO). Experiments were performed in accordance with NIH Guidelines for the Care and Use of Laboratory Animals, and all procedures were approved by the Institutional Animal Care and Use Committee of the University of Southern California.

### 2.2. Stereotaxic cannula implantation

Under 2-4% isoflurane delivered at a rate of 250 mL/min, rats were surgically implanted with either unilateral or bilateral 26 G guide cannulae (Plastics One, Roanoke, VA) targeting either the lateral ventricle (LV): −0.9 mm anterior/posterior (AP), +1.8 mm medial/lateral (ML), −2.6 mm dorsal/ventral (DV) (reference point for AP and ML at bregma, reference point for DV at skull surface at target site), or nucleus accumbens shell (ACBsh): +1.1 mm anterior/posterior (AP), ±1.0 mm medial/lateral (ML), −5.1 mm dorsal/ventral (DV), respectively.

Placement for the LV cannula was verified by elevation of cytoglucopenia resulting from an injection of 210 μg (2μl) of 5-thio-D-glucose (5tg) (Slusser and Ritter, 1980) using an injector that extended 2mm beyond the end of the guide cannula. A post-injection elevation of at least 100% of baseline glycemia was required for subject inclusion. Animals that did not pass the 5tg test were retested with an injector that extended 2.5 mm beyond the end of the guide cannula and, upon passing 5tg, were subsequently injected using a 2.5-mm injector instead of a 2-mm injector for the remainder of the study.

For targeting the ACBsh, injectors (33G; Plastics One, Roanoke, VA) extending 2.0 mm below the end of the guide cannulas were used. ACBsh cannula placement was verified at the conclusion of experimental testing via histological identification of bilateral 100nl 2% pontamine sky blue ink (2%) injection sites based on the Swanson Rat Brain Atlas (Swanson, 2003). Injection sites within the boundaries of the ACBsh were considered correct, and only data from rats with correct placement were included in the analyses (89% hit rate). Photomicrographs were acquired using a Nikon 80i (Nikon DSQI1,1280X1024 resolution, 1.45 megapixel) under darkfield condenser (Representative injection site, Fig. 1A).

**Figure 1:**
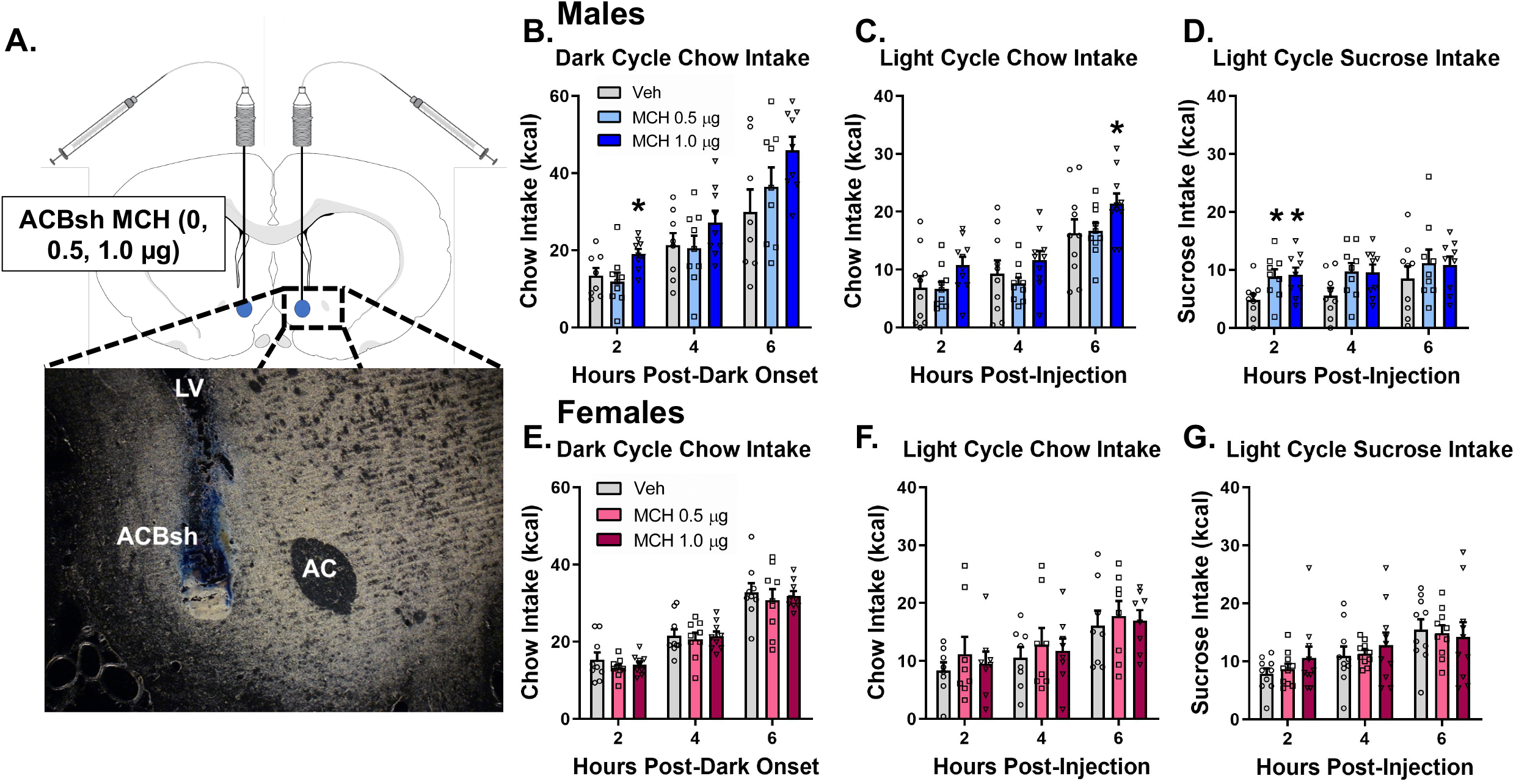
Pharmacological activation of ACBsh MCH1R promotes feeding in a sex-specific manner. A representative ACBsh injection site is shown as localization of pontamine sky blue ink following a 100 nl injection (a). Direct activation of MCH1R in the ACBsh MCH significantly increased cumulative dark cycle chow intake (b), light cycle chow intake (c), and light cycle sucrose intake (d) in male rats. ACBsh MCH did not affect food intake during the same feeding tests in female rats (e-g). Data are mean ± SEM; *p < 0.05 vs vehicle treatment.

### 2.3. Intracranial virus injections

For stereotaxic injections of viruses and tracers, rats were first anesthetized using 2-4% isoflurane delivered at a rate of 250 mL/min. Viruses were delivered using a microinfusion pump (Harvard Apparatus, Cambridge, MA, USA) connected to a 33-gauge microsyringe injector attached to a PE20 catheter and Hamilton syringe. Flow rate was calibrated and set to 5 μl/min; injection volume was 200 nl/site. Injectors were left in place for 2 min post-injection. Following AAV injections, animals were surgically implanted with a unilateral cannula targeting the lateral ventricle using the coordinates listed above. All experimental procedures occurred 21 days post virus injection to allow for transduction and expression. Successful virally mediated transduction was confirmed postmortem based on the Swanson Brain Atlas (Swanson, 2003) in all animals via IHC staining using immunofluorescence-based antibody amplification to enhance the fluorescence transgene signal, followed by manual quantification under epifluorescence illumination using a Nikon 80i (Nikon DS QI1,1280X1024 resolution, 1.45 megapixel).

For the dual-virus approach for selective expression of DREADDs in MCH neurons that project to the ACBsh, bilateral injections of the AAV2(retro)-eSYN-EGFP-T2a-icre-WPRE (AAV2-retro CRE; 200 nl; titer 1.7×10^13^ GC/ml) were delivered to the ACBsh using the following coordinates: +1.1 mm AP, ±0.8 mm ML, −7.5 mm DV (from the skull at bregma). Bilateral stereotaxic injections where then given for the AAV2-DIO-rMCHp-hM3D(Gq)-mCherry (Cre-dependent MCH DREADDs; titer 1.1×10^13^ GC/ml) at the following coordinates: injection (1) −2.6 mm AP, ±1.8 mm ML, −8.0 mm DV; (2) −2.6 mm AP, ±1.0 mm ML, −8.0 mm DV; (3) −2.9 mm AP, ±1.1 mm ML, −8.8 mm DV; (4) −2.9 mm AP, ±1.6 mm ML, −8.8 mm DV (from the skull surface at bregma). Our previous publications show that this AAV is selective to MCH neurons (IHC verification), and that DREADDs are functional in MCH neurons (behavioral and electrophysiological verification) (Noble et al., 2018; Noble et al., 2019). Present results are comparable to our previous application of this method in the ACBsh, which yielded ~10% of total MCH neurons transfected with DREADDS, and ~100% of cells transfected with DREADDs that were also MCH+ neurons (Noble et al., 2018). Moreover, using this approach, ICV CNO-mediated activation of MCH neurons increased chow intake, and these effects were blocked by pretreatment with the MCH1R antagonist, H6408 (10 μg) (Noble et al., 2018).

### 2.4. Immunohistochemistry

Rats were anesthetized and sedated with a ketamine (90 mg/kg)/xylazine (2.8 mg/kg)/acepromazine (0.72 mg/kg) cocktail, then transcardially perfused with 0.9% sterile saline (pH 7.4) followed by 4% paraformaldehyde (PFA) in 0.1M borate buffer (pH 9.5; PFA). Brains were dissected out and post-fixed in PFA with 12% sucrose for 24 h, then flash frozen in isopentane cooled in dry ice. Brains were sectioned to 30-μm thickness on a freezing microtome. Sections were collected in 5 series and stored in antifreeze solution at −20°C until further processing. General fluorescence IHC labeling procedures were performed, as reported (Noble et al., 2018; Suarez et al., 2018). The following antibodies and dilutions were used: rabbit anti-RFP (1:2000, Rockland Inc., Limerick, PA, USA) and rabbit anti-MCH (1:1000; Phoenix Pharmaceuticals, Burlingame, CA, USA). Antibodies were prepared in 0.02M potassium phosphate buffered saline (KPBS) solution containing 0.2% donkey serum and 0.3% Triton X-100 at 4 °C overnight. After thorough washing with 0.02M KPBS, sections were incubated in secondary antibody solution. Secondary antibodies were obtained from Jackson Immunoresearch and used at 1:500 dilution at 4 °C, with overnight incubations (Jackson Immunoresearch; West Grove, PA, USA). Sections were mounted and coverslipped using 50% glycerol in 0.02M KPBS and the edges were sealed with clear nail polish. IHC detection of Photomicrographs were acquired using a Nikon 80i (Nikon DSQI1,1280X1024 resolution, 1.45 megapixel) under epifluorescence.

### 2.5. Characterization of DREADDs expression

DREADDs expression was quantified in 1 out of 5 series of brain tissue sections from the perfused brains cut at 30 μm on a freezing microtome based on counts for the fluorescence transgene, mCherry. Immunofluorescence staining for red fluorescent protein (RFP) was conducted as described above to amplify the mCherry signal. Counts were performed in sections from Swanson Brain Atlas level 27–32 (Swanson, 2003), which encompasses all MCH-containing neurons (Hahn, 2010). Cell counts were performed in all dual-virus Cre-dependent MCH DREADDs-injected animals, and animals with 80 or fewer RFP positive cells (a priori established inclusion criterion) were excluded from all experimental analyses. Counts were performed using epifluorescence illumination using a Nikon 80i (Nikon DS-QI1,1280X024 resolution, 1.45 megapixel).

### 2.6. Drug preparation and intracranial pharmacological injections

For ACBsh injections of MCH, MCH (Bachem Americas, Torrance, CA, USC; Cat #: H-2218.1000) was dissolved in aCSF and diluted to 5 μg/μl or 2.5 μg/μl. For chemogenetic activation of MCH neurons, CNO (National Institute of Mental Health; 18 mmoles in 2 μl) or 33% dimethyl sulfoxide (DMSO) in aCSF (daCSF; vehicle control in 2 μl) was administered ICV. Animals were handled and habituated to injections prior to testing. All injections were delivered through a 33-gauge micro-syringe injector attached to a PE20 catheter and Hamilton syringe. For ACBsh injections, a microinfusion pump (Harvard Apparatus) was used. The flow rate was set to 5 μl/min and 100 nl bilateral injection volume. Injectors were left in place for 30 s to allow for complete infusion of the drug. For ICV injections, 2 μl (CNO and daCSF) was delivered by manually via a Hamilton syringe.

### 2.7. Food intake studies

Food intake analyses occurred in the animal’s home cage. For both studies using laboratory chow (LabDiet 5001, LabDiet, St. Louis, MO, USA) and 45-mg sucrose pellets (TestDiet, Richmond, IN, USA) home cage food was removed 2 h prior to start of the intake analyses.

Animals were randomized to receive either aCSF, 0.5 μg, or 1 μg MCH using a counterbalanced within-subjects design. Injections were given 45 min prior to the start of the test and pre-weighed amounts of the test diet were deposited in the home cage immediately after the lights went out (for dark cycle chow intake studies, male (n = 9) and female (n=10) rats) or during the early midlight phase (4 h post-light onset) (for light cycle intake studies). Sample sizes for light cycle chow intake were male (n =10) and female (n = 8) rats; sample sizes for light cycle sucrose were male (n = 9) and female (n = 10) rats.

For chemogenetic activation of MCH neurons, a counterbalanced, within-subjects design was used such that rats receive ICV injection of either CNO (National Institute of Mental Health; 18 mmol in 2 μl) or 33% DMSO in aCSF (daCSF; vehicle control in 2 μl). Injections were given 1 h prior to the lights going off and pre-weighed amounts of the test diet were deposited in the home cage immediately after the lights went out (for dark chow cycle intake studies; male (n = 10) and female (n=10) rats) or 4 h post-light onset (for light cycle intake studies). Sample sizes for both light cycle chow and light cycle sucrose intake were male (n = 10) and female (n = 11) rats. A control experiment was also conducted in male rats to examine possible effects of ICV CNO on food intake [daCSF (n = 8) and CNO (n=8)]. Spill papers were placed underneath the cages to collect food spillage, which was weighed and added to the difference between the initial hopper weight and the hopper weight at each measurement time point. A total of 72 h was allotted between treatments.

### 2.8. PROG sucrose/chow choice test

Using the modified version of the protocol described in (Randall et al., 2012), male rats (n = 11) were chronically food restricted to maintain 85% free-feeding body weight, and then trained to press an active lever for 45-mg sucrose pellets, starting with one lever press to one sucrose pellet (FR1). After three weeks of training, rats were then trained on a progressive (PROG) procedure. For PROG sessions, the ratio started at FR1and was increased by one additional response every time 15 reinforcements were obtained (FR1 x 15, FR2 x 15, FR3 x 15, FR4 x 15, …). After one week of PROG training, free access to chow was introduced concurrently and available during PROG sessions for one week of training. All training and testing sessions were 30 minutes. Following this training period, a counterbalanced, within-subjects design was used to assess the effect of MCH in the ACBsh on performance on the PROG vs chow task. Rats were injected with either aCSF or 1 μg MCH into the ACBsh 45 minutes prior to the start of the task. The ratio of lever presses to sucrose pellets earned (PROG ratio) and free chow intake was recorded. A total of 72 hours was allotted between treatments. All training and tests occurred during the dark cycle.

### 2.9. Open field test

The apparatus used for the open field test is an opaque gray plastic bin (60 cm × 56 cm), which was positioned on a flat table in an isolated room with a camera directly above the center of the apparatus. Desk lamps were positioned to deliver indirect light on all corners of the maze such that the lighting in the box measured 30 lux in all corners and in the center. At the start of the 10-min test, each rat, male (n = 10) and female (n = 11), is placed in the open field apparatus in the same corner facing the center of the maze. All sessions were video recorded and ANY-maze video tracking software (Stoelting, Wood Dale, IL) was used for activity tracking. Total distance traveled was measured by tracking movement from the center of the rat’s body. On test days, food was removed 2 h prior to the lights going off and behavioral testing began when the lights turned off. For investigating the effects of stimulating ACBsh-projecting MCH neurons on open field activity levels, animals were randomized to receive either intra-LV daCSF or CNO using a counterbalanced within-subjects design, with a washout period of 72 h between the two treatments. Injections were given 45 min prior to behavioral testing.

### 2.10. Ovariectomy surgery

In order to determine whether estradiol modulates the orexigenic response to MCH, female rats underwent bilateral ovariectomy using an intra-abdominal approach under 2-4% isoflurane delivered at a rate of 250 mL/min. Briefly, an abdominal incision was made, and the uterus was located. The fallopian tube on one side was coaxed out until the ovary was visible. When enough of the fat pad holding the ovaries was visible, an autoclaved suture was used to tightly tie off the tube about mid-way between ovary and uterus. The cautery tool was then used to separate the ovary from the fat pad. The fallopian tube was cut between the ovary and the suture that was placed to fully remove the ovary. This was then repeated on the other ovary.

### 2.11. Estrogen replacement and food intake study

β-Estradiol-3-benzoate (EB) (Sigma-Aldrich, St. Louis, MO) was dissolved in sesame oil. One week following surgery, rats were separated into weight-matched groups and placed on a cyclic regimen of either 2 μg β-estradiol-3-benzoate (EB) delivered in 0.1ml sesame oil or oil vehicle alone (EB (n = 9) and oil (n = 8) rats). This dose of EB was chosen because it has been shown to produce plasma estradiol levels comparable to that which is achieved physiologically during proestrus in intact cycling rats (Asarian and Geary, 1999). Furthermore, this dose has been shown to induce a transient decrease in food intake, similar to the decrease observed during estrus in cycling rats (Asarian and Geary, 2002). To assess EB’s ability to affect the orexigenic effect of MCH in the ACBsh, we conducted feeding tests the day after rats received their weekly estradiol or oil injection, allowing us to examine EB’s effect at a time during which EB-induced suppression of food intake is expressed. The EB treatment timing of every 4-5 days has been previously shown to be an effective regimen (Asarian and Geary, 1999; Geary and Asarian, 1999; Geary et al., 1994; Santollo and Eckel, 2008a, b). This regimen mimics the changes in plasma estradiol levels across the ovarian cycle in gonadally intact females (Asarian and Geary, 2002). That is, estradiol levels are low, then peak on the day of EB treatment, and rapidly fall the subsequent day, which models estrus (Becker et al., 2005). A within-subjects counterbalanced design was used to assess the feeding response to intra-ACBsh MCH in oil- and EB-treated rats. On test days, chow was removed 2 h prior to start of the intake analyses. Rats received intra-ACBsh aCSF or 1.0 μg MCH 45 minutes before chow was returned at 4 h post-light onset (early midlight phase) and food intake was subsequently measured.

### 2.12. Fluorescence in situ hybridization

Tissue sections from male (n = 3) and female (n = 3) rats were mounted on subbed glass slides (Fisher brand Superfrost Plus, Fisher Scientific, Hampton, NH, USA) and desiccated overnight (~16 h) in a vacuum desiccant chamber. Following 1 h and 45 min postfix in 4% PFA, sections were washed 5 × 5 min in KPBS and incubated for 30 min at 37 °C in a solution of 100mM Tris (pH 8), 50mM EDTA (pH 8), and 0.1% Proteinase K (10 mg/ml, Sigma P2308), then rinsed for 3 min in the same Tris and EDTA solution without Proteinase K. Sections were washed 3 min in a solution of 100mM triethanolamine (pH 8) in water and then incubated for 10 min at room temperature with 0.25% acetic anhydride in 100mM triethanolamine, then washed 2 × 2 min in 10% 20× saline-sodium citrate buffer. Prior to hybridization, sections were dehydrated in increasing concentrations of ethanol (50%, 70%, 95%, 100%, 100%). Sections were incubated with probes for 3 h (MCHR1 and ESR1 probes). Reagents from the RNAscope^®^ Fluorescent Multiplex Detection Reagent Kit v2 (Advanced Cell Diagnostics, Newark, CA, USA) were used to amplify the probe as per the kit’s instructions. Slides were coverslipped using ProLong^®^ Gold Antifade Reagent (Cell Signaling, Danvers, MA, USA; Cat #: 9071s). Photomicrographs were acquired using a Nikon 80i (Nikon DS-QI1,1280X1024 resolution, 1.45 megapixel) under epifluorescent illumination using the Nikon Elements BR software.

### 2.13. Statistical analyses

Data are reported as means ± SE. For all behavioral experiments, repeated measures ANOVA and post hoc Holm-Bonferroni comparisons were made. Alpha levels were set to α = 0.05 for all analyses.

## 3. Results

### 3.1. Pharmacological activation of ACBsh MCH1R promotes feeding in a sex-specific manner

To determine the effects of ACBsh MCH1R stimulation on dark cycle chow intake, male rats received bilateral intra-ACBsh injection of vehicle or MCH at doses subthreshold for effect when delivered to the ventricle (0.5 or 1.0 μg) (Noble et al., 2019) 45 min prior to dark onset. ACBsh MCH1R activation significantly increased dark cycle chow intake, as evident from a significant main effect of ACBsh MCH on cumulative chow intake at 2 h [F(2,16) = 8.188, *P* < 0.01] post-dark onset. Post hoc analysis revealed the higher dose,1.0 μg, increased intake at 2 h post-dark onset relative to aCSF (*P* < 0.01) (Fig. 1B). Given that separate one-way ANOVAs of cumulative intake at each time point can potentially increase false positives, we also include additional analyses of noncumulative 2hr binned intake data with time as a repeated measures factor (Fitts, 2006). These noncumulative 2hr binned analyses revealed a trend toward both a significant time x drug interaction [F(4,32) = 2.57, *P* = 0.057] and main effect of drug [F(2,16) = 2.97, *P* = 0.080] (Supplemental Fig. 1A). We also examined the effect of ACBsh MCH1R activation on light cycle chow intake in male rats. ACBsh MCH tended to increase cumulative light cycle chow intake a 2 h post-injection [F(2,18) = 3.115, *P* = 0.07] and significantly increased light cycle chow intake at 6 h post-injection [F(2,18) = 4.287, *P* < 0.05]. Post hoc comparisons revealed that at 6 h post-injection, 1.0 μg of MCH in the ACBsh significantly increased chow intake relative to aCSF (*P* < 0.05) (Fig 1C). Noncumulative 2hr binned analysis of light cycle chow intake in males revealed a trend toward a significant time x drug interaction [F(4,36) = 1.86, *P* = 0.10] and significant main effect of drug [F(2,18) = 4.287, *P* < 0.05] (Supplemental Fig. 1B). We next determined the effect of ACBsh MCH1R stimulation on light cycle sucrose intake in male rats. ACBsh MCH significantly increased cumulative sucrose intake at 2 h post-injection [F(2,16) = 3.911, *P* < 0.05] with a trend observed at 4 h post-injection [F(2,16) = 3.083, *P* = 0.08]. Post hoc comparisons showed that both the low and high dose of MCH increased sucrose intake at the 2 h timepoint relative to aCSF (*P* < 0.05 for both) (Fig 1D). Noncumulative 2hr binned analysis of light cycle sucrose intake revealed a significant time x drug interaction [F(4,32) = 3.294, P < 0.05] with the effect of ACBsh MCH to increase sucrose intake occurring during the 0-2hr time bin (Supplemental Fig. 1C).

The same food intake analyses were conducted in female rats. Results revealed no significant effects of intra-ACBsh MCH injections at any timepoint for cumulative dark onset chow intake, light cycle chow intake, or sucrose intake measured in light cycle in female rats

(Fig 1E-G). Moreover, noncumulative 2hr binned feeding analysis reveal no significant effects of ACBsh MCH. Two-way repeated measures ANOVAs revealed a significant main effect of time, however there was no main effect of drug and no time x drug interaction for noncumulative 2hr binned dark onset chow intake, light cycle chow intake, or sucrose intake in female rats (Supplemental Fig. 1D-F).

### 3.2. Chemogenetic stimulation of ACBsh-projecting MCH neurons promotes food intake in a sex-specific manner

Here we used a dual virus chemogenetic approach to bilaterally activate ACBsh-projecting MCH neurons (Fig 2A). Consistent with our previous report using this approach (Noble et al., 2018), histological analyses confirmed that all tdTomato+ neurons coexpress MCH, and that approximately ~10% of all MCH neurons are tdTomato+ with AAV Retro-Cre targeted to the ACB (Fig 2B). In order to determine the effect of activating ACBsh-projecting MCH neurons on dark cycle chow intake, male rats received LV injection of vehicle or 18 mmol clozapine-N Oxide (CNO; DREADDs ligand) 1 hour prior to dark onset. ICV CNO significantly increased cumulative dark cycle chow intake at 2 h [t(9) = 2.416, *P* < 0.05] post-dark onset (Fig 3A). Noncumulative 2hr binned analysis of dark cycle chow intake revealed a significant time x drug interaction [F(2,18) = 6.259, P < 0.01]. Post hoc comparisons showed a significant increase in noncumulative chow intake during the 0-2 h time bin and a significant reduction in feeding during the 2-4 h time bin following CNO (Supplemental Fig 2A). We next examined the effect of stimulation of ACBsh-projecting MCH neurons on light cycle chow intake in male rats. During the light cycle, CNO injection significantly increased cumulative chow intake at 2 h [t(9) = 1.916, *P* < 0.05] and 4 h [t(9) = 2.725, *P* < 0.05] post-injection relative to vehicle (Fig 3B). Noncumulative 2hr binned analysis of light cycle chow intake in males revealed a trends toward a significant time x drug interaction [F(2,18) = 2.25, *P* = 0.13] and main effect of drug [F(1,9) = 2.945, *P* = 0.13] (Supplemental Fig 2B). In order to determine the role of stimulating ACBsh-projecting MCH neurons on intake of a palatable food, male rats received ICV CNO or vehicle and were subsequently given 6-h access to sucrose pellets in the home cage during the light cycle. Here we found that CNO significantly increased cumulative sucrose intake at 6 h post-injection relative to vehicle [t(9) = 2.172, *P* < 0.05] (Fig 3C). Noncumulative 2hr binned analysis of light cycle sucrose intake in males revealed a trend toward a significant main effect of drug [F(1,9) = 4.71, *P* = 0.58] (Supplemental Fig 2C). As a control experiment for possible effects of CNO independent of DREADDs (Manvich et al., 2018), male rats that did not receive dual viral injections received ICV vehicle or 18 mmol CNO 1 hour prior to dark onset. ICV CNO did not affect either cumulative and noncumulative 2hr binned dark cycle chow intake in these rats (Supplemental Fig 3).

**Figure 2:**
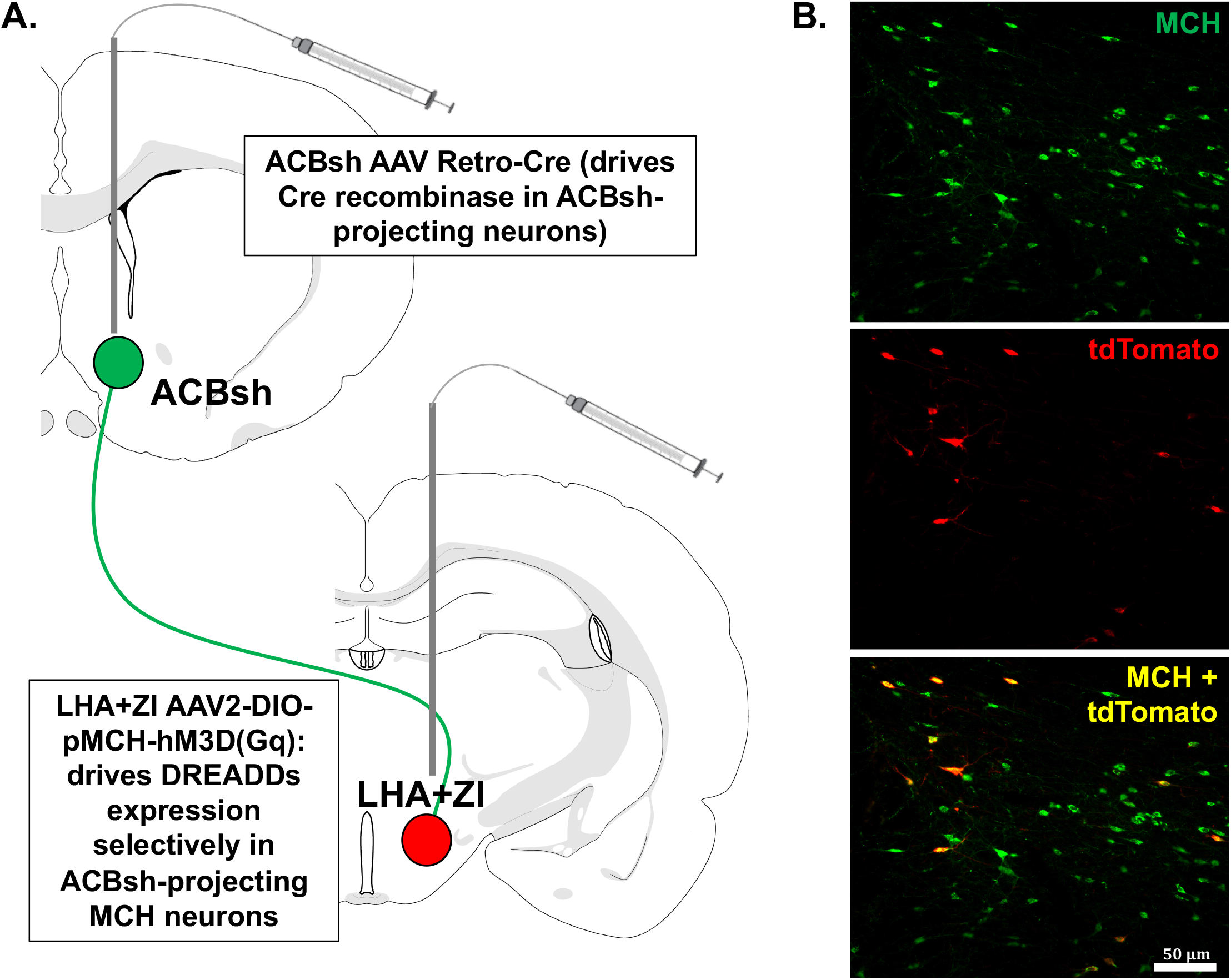
A schematic diagram representing the dual-virus approach for selective expression of hM3D(Gq) DREADDs in MCH neurons that project to the ACBsh. Bilateral injections of the AAV2(retro)-eSYN-EGFP-T2a-icre-WPRE were delivered to the ACBsh to drive Cre recombinase in ACBsh-projecting neurons. Bilateral stereotaxic injections where then delivered to the LHA and ZI for the AAV2-DIO-rMCHp-hM3D(Gq)-mCherry (Cre-dependent MCH DREADDs) to drive DREADDs expression selectively in ACBsh-projecting MCH neuron (a). Representative images showing the localization of the mCherry fluorescence reporter (tdTomato) in MCH neurons (green) following dual-virus injections (colocalization in yellow) (b).

**Figure 3:**
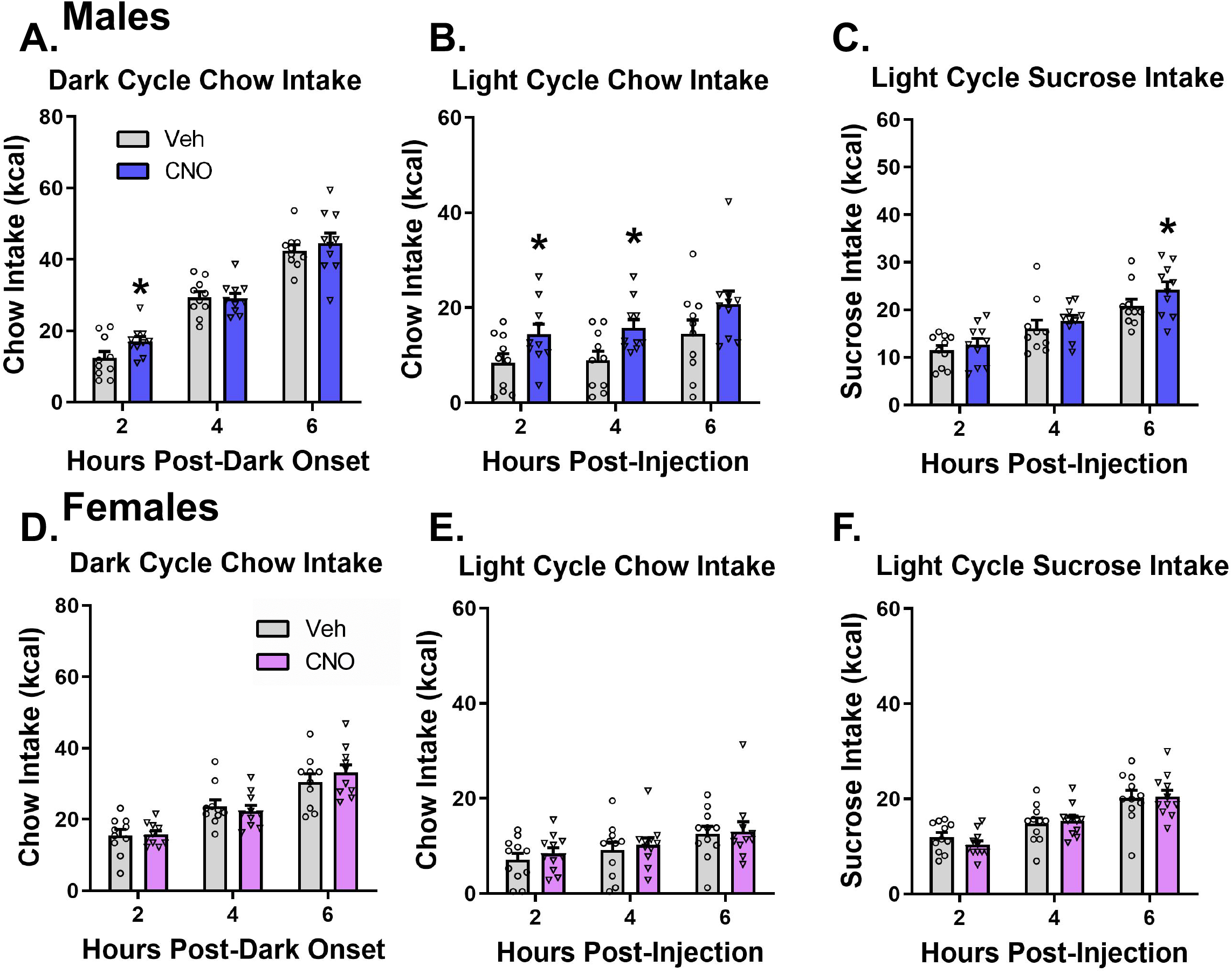
Chemogenetic stimulation of ACBsh-projecting MCH neurons promotes food intake in a sex-specific manner. In male rats, activation of ACBsh-projecting MCH neurons following ICV CNO significantly increased cumulative dark cycle chow intake (a), light cycle chow intake (b), and light cycle sucrose intake (c). Activation of ACBsh-projecting MCH neurons had no effect on cumulative dark cycle chow intake, light cycle chow intake, or light cycle sucrose intake in female rats (d-f). Data are mean ± SEM; *p < 0.05 vs vehicle treatment.

The same food intake analyses were conducted separately in female rats. Activation of ACBsh-projecting MCH neurons had no effect on either cumulative (Fig 3D-F) or noncumulative 2hr binned (Supplemental Fig 2D-F) dark cycle chow intake, light cycle chow intake, or light cycle sucrose intake at any timepoint in female rats.

### 3.3. Activation of ACBsh MCH1R does not influence effort-based food choice

The ACBsh is well established as an important site for incentive motivational processes. To investigate whether ACBsh MCH signaling affects choice behavior between effort-based palatable food vs. no/low effort consumption of bland food, food-restricted male rats were trained to lever press for a 45 mg sucrose reinforcer on a PROG schedule with free chow also made available during the sessions. Results from within-subjects Student’s t-test reveal that intra-ACB MCH had no significant effect on pellets earned, PROG ratio, chow intake, or sucrose ratio during the PROG vs. chow choice task (Fig 4B-E). Furthermore, there were no differences in presses on the non-reinforced lever (Student’s two-tailed, paired t test; means [SEM]: Vehicle 2.18 [1.3], MCH 1.09 [0.5]; n = 11).

**Figure 4:**
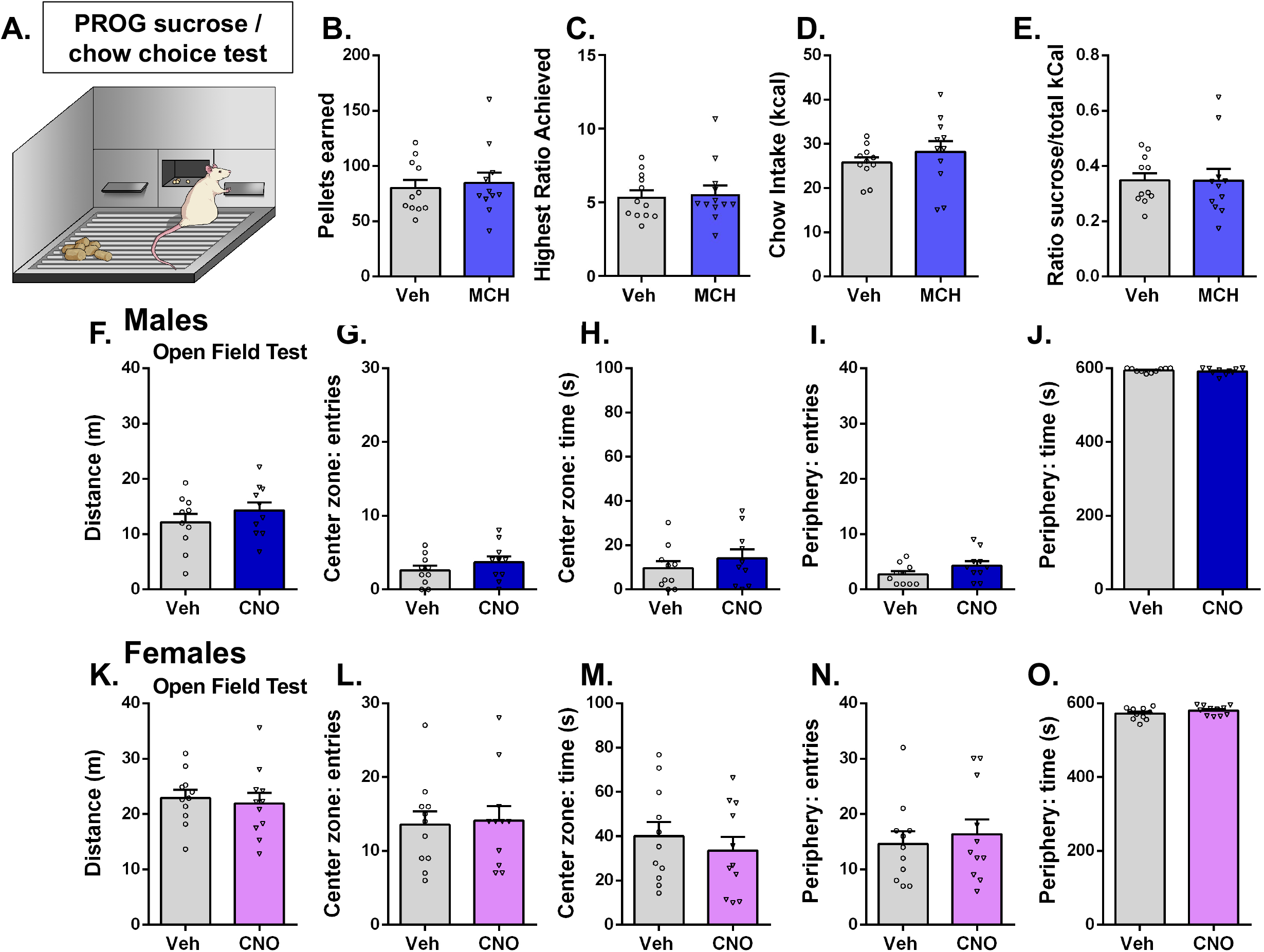
Activation of ACBsh MCH1R does not influence effort-based food choice (high effort sucrose vs. low/no effort chow). The same dose of MCH (1.0 μg) that significantly increases feeding in the home cage had no significant effect on pellets earned (b), PROG ratio (c), chow intake (d), or sucrose ratio (e) during the PROG vs. chow choice task in male rats. Chemogenetic stimulation of ACBsh-projecting MCH neurons does not influence locomotor or anxiety-like behavior. In both sexes, ICV CNO did not influence distance traveled, center zone entries, center zone time, periphery entries, or periphery time. (Males: f-j; Females: k-o). Data are mean ± SEM.

### 3.4. Chemogenetic stimulation of ACBsh-projecting MCH neurons does not influence locomotor or anxiety-like behavior

ACB neural processing is associated with locomotor activity and anxiety-like behavior (Barrot et al., 2005; Chee et al., 2019; Pijnenburg and Rossum, 1973; Yamada et al., 2020; Zhu et al., 2016). In order to test whether activation of ACBsh-projecting MCH cells influences general levels of physical activity or anxiety-like behavior, we utilized an open field test. Results from within-subjects Student’s t-test for each variable reveal that in both male and female rats, ICV CNO treatment did not significantly influence distance traveled during the test session (Males: Fig 4F, Females: Fig 4K), nor did CNO significantly influence center zone entries, center zone time, periphery entries, or periphery time, measures of anxiety-like behavior (Males: Fig 4G-J, Females: Fig 4L-O).

### 3.5. Estradiol reduces the orexigenic response of ACBsh MCH signaling

Based on our findings that stimulation of MCH1R in the ACBsh increased food intake in males but not in intact cycling females, we hypothesized that the orexigenic effect of intra-ACBsh MCH may be attenuated by estradiol. Two-way mixed-design ANOVA at each timepoint reveal a significant main effect of MCH to increase cumulative light cycle chow intake at 2 h [F(1,15) = 5.387, *P* < 0.05], 4 h [F(1,15) = 6.706, *P* < 0.05], and 6 h [F(1,15) = 7.728, *P* < 0.05] post-injection. Pairwise planned comparisons revealed that in the oil-treated rats, 1.0 μg of MCH delivered to the ACBsh significantly increased cumulative chow intake at 2 and 6 h post-injection (*P* < 0.05 for both) (Fig 5A). Whereas in the EB-treated rats, no significant effects of ACBsh MCH injections were observed at any timepoint. Conclusions from these planned comparisons should be taken with caution as the hormone x drug interaction did not achieve significance at any time point (*Ps* > 0.05). Noncumulative 2hr binned analysis of light cycle chow intake with three-way mixed-design ANOVA revealed both a significant main effect of time [F(2,30) = 33.43, P < 0.0001] and a main effect of drug [F(1,15) = 7.66, P < 0.05]. Two-way mixed-design ANOVA revealed a significant main effect of MCH to increase noncumulative light cycle chow intake only at 0-2 h [F(1,15) = 5.387, *P* < 0.05] (Supplemental Fig 4).

**Figure 5:**
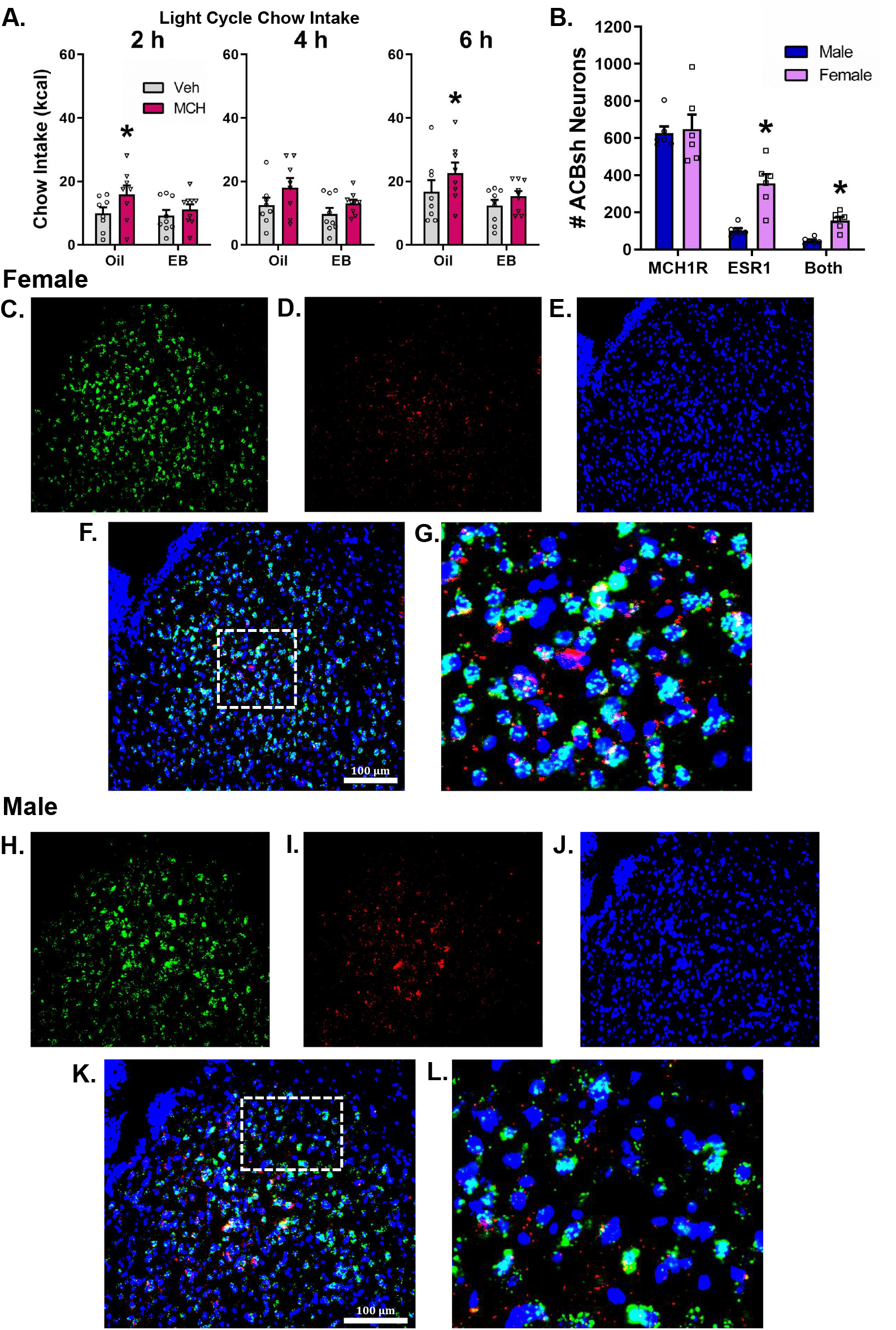
Estradiol reduces the orexigenic response of ACBsh MCH signaling. In the oil-treated OVX female rats, 1.0 μg of MCH delivered to the ACBsh significantly increases cumulative light cycle chow intake; whereas in the EB-treated OVX rats, ACBsh MCH injections did not affect chow intake (a). There was no sex difference in the number of MCH1R positive neurons in the ACB. In female rats, neurons in the ACBsh contain significantly more overall ESR1 mRNA relative to males, and females had significantly more neurons in the ACBsh that contained both MCH1R and ESR1 relative to male rats (b). Data are mean ± SEM; *p < 0.05. Representative images for both females and males showing fluorescence in situ hybridization for MCHR1 mRNA (green) (c,h) and the ESR1 mRNA (red) (d,i), with DAPI counterstain (blue) (e,j) in the ACBsh. Merged representative images show that MCH1R and ESR1 mRNA expression co-localize in the ACBsh to a greater extent in females (Males: k,l; Females: f,g).

### 3.6. MCH1R and ESR1 mRNA expression co-localize in the ACBsh to a greater extent in females

We observed no difference in the number of MCH1R positive neurons in the ACBsh in male and female rats, (Fig 5B). However, we found that in female rats, neurons in the ACBsh contain significantly more overall ESR1 expression relative to males. Furthermore, female rats had significantly more neurons in the ACBsh that contained both MCH1R and ESR1 relative to male rats. We found that in males, 8% of MCH1R containing neurons in the ACBsh also contain ESR1, whereas in females, ~25% of MCH1R containing neurons in the ACBsh also contain ESR1. Moreover, in both male and female rats, ~45% of ESR1 containing neurons in the ACBsh also contain MCH1R.

## 4. Discussion

MCH is a neuropeptide produced in the lateral hypothalamic area (LHA) and the adjacent zona incerta (ZI) that potently increases food intake via unidentified neural pathways. Here we show that MCH signaling in the ACBsh, a region embedded in the central reward system, potently increases food intake in a sex-dependent manner. Both site-specific pharmacological activation of ACBsh MCH1R and chemogenetic-mediated activation of ACBsh-projecting MCH neurons increased intake of standard chow as well as palatable sucrose in male rats. However, these same approaches did not affect food intake in intact cycling female rats, suggesting that the ability of MCH in the ACBsh to increase feeding may be dependent on endogenous female reproductive hormones. In order to directly test this hypothesis, we examined chow intake following ACBsh MCH1R stimulation in ovariectomized (OVX) female rats on a cyclic regimen of either 2 μg β-estradiol-3-benzoate (EB) or oil vehicle. We found that ACBsh MCH1R stimulation significantly increased chow intake only in oil-treated rats, suggesting that estradiol attenuates orexigenic effect of ACBsh MCH signaling. Consistent with the sex and estradiol influences on ACBsh MCH-mediated feeding effects, fluorescent in situ hybridization analyses confirm that MCH1R and estrogen receptor 1 (ESR1) mRNA expression co-localize in the ACBsh to a greater extent in females vs. males. Collective results reveal that that MCH-ACBsh signaling promotes eating in an estradiol- and sex-dependent manner, thus identifying novel neurobiological mechanisms through which MCH and estrogen influence food intake.

ACB MCH1R-expressing neurons and glial cells are not exclusively engaged by MCH signaling under physiological conditions, but also by various neurochemicals that are co-released by LHA/ZI MCH neurons. Here we utilized our recently validated dual-viral chemogenetic approach (Noble et al., 2018; Noble et al., 2019) to reversibly and selectively activate MCH neurons that project to the ACBsh. Similar to ACBsh-targeted MCH1R pharmacology, results reveal that ACBsh-projecting MCH neuron activation increases intake of both chow and palatable sucrose in male rats, thus revealing a novel role for this population of neurons in feeding behavior. Given that the hyperphagic effects of activating ACBsh-projecting MCH neurons were similar to those following ACBsh MCH injections, it is likely that MCH is a key signaling molecule mediating the effects of MCH neurons on eating. This notion is further supported by data showing that the increased chow intake following chemogenetic-mediated activation of a heterogenous population of MCH neurons (including those projecting to the ACBsh) are completely blocked by pretreatment with the MCH1R antagonist, H6408 (Noble et al., 2018). One important additional consideration with the chemogentic approach is the branching collateral projections of single neurons to more than one target brain region. Indeed, MCH neurons have extensive projections throughout the neuraxis and very little is understood about the collateral targets of specific MCH projection pathways. It is therefore possible that MCH neurons that project to the ACBsh may also project to other brain regions, and that observed results may be based, in part, on these collateral projection targets. However, given the comparable magnitude and timeframe of the hyperphagic effects of pharmacological ACBsh MCH administration and chemogenetic activation of ACBsh-projecting MCH neurons, collective results suggest that MCH collateral signaling to other brain regions is unlikely to be mediating the observed feeding outcomes following ACBsh-targeted MCH neuron activation.

The ACBsh plays a critical role in reward-motivated behaviors and increased ACB dopamine signaling alters food choice behavior in the PROG sucrose/chow choice test in favor of high-effort sucrose compared to low/no effort bland chow (Randall et al., 2012). Here we found that the same dose of MCH in the ACBsh that increased free consumption of both chow and sucrose within the first two hours after injections in the home cage did not affect food choice behavior in the PROG test, thus suggesting that increased incentive motivation for sucrose pellets is unlikely to underlie the observed hyperphagia associated with ACBsh MCH1R stimulation. These findings also suggest that the orexigenic effects of ACBsh MCH signaling are unlikely based on elevated dopamine signaling. In fact, previous work suggests that MCH signaling reduces ACB dopamine signaling, as pharmacological activation of ACB MCH1R blocks dopamine-induced glutamatergic receptor phosphorylation (Georgescu et al., 2005), and MCH1R deletion from ACB GABAergic neurons is associated with a hyperdopaminergic state (Chee et al., 2019). In contrast, optogenetic activation of MCH neurons (nonspecific with regards to projection target) increases ACB dopamine release when activation is paired with consumption of the nonnutritive sweetener sucralose (Domingos et al., 2013). While more research is needed to fully understand the influence of MCH signaling on striatal dopamine responses, present results indicate that ACBsh MCH is unlikely to increase feeding behavior based on increased dopamine signaling and/or increased incentive salience for palatable food reinforcement. Future studies are also necessary to determine whether ACBsh MCH1R signaling influences motivated responding for sucrose under different testing parameters, and whether chemogenetic activation of ACBsh-projecting MCH neurons plays a role in motivation for sucrose.

Recent data show that, in addition to producing a hyperdopaminergic state, MCH1R deletion from ACB GABAergic neurons in mice leads to increased locomotor activity (Chee et al., 2019). Further, systemic pharmacologic MCH1R blockade in mice is associated with hyperactivity (Zhang et al., 2014). Thus, it is possible that our observed ACBsh MCH-driven increases in food intake are secondary to altered physical activity levels. However, selective chemogenetic activation of ACBsh-projecting MCH neurons had no effect on activity levels in the open field test, thus suggesting that altered activity levels are unlikely to be causally related to the observed orexigenic effects in the present study. It is possible that the contrast between the lack of physical activity effects in the present study vs. the increased locomotor activity observed following MCH deletion from ACB GABA neurons is based on extra ACBsh MCH1R signaling (ACB core) in this latter study, and/or species (rats vs. mice) or methodological (chemogenetic vs. transgenic deletion) differences.

Previous studies demonstrate that MCH signaling is also involved in depression-like behavior (Borowsky et al., 2002; Lagos et al., 2011; Takekawa et al., 2002; Ye et al., 2018), and that these effects may involve ACB MCH1R signaling. For example, blockade of MCH1R in the ACBsh produces an antidepressant-like effect in the forced swim test, whereas ACBsh injection of MCH has the opposite effect (Georgescu et al., 2005). While we did not directly examine depression-like behavior in the present study, our results from the open field test demonstrate that anxiety-like behavior, measured from center zone entries, center zone time, periphery entries, or periphery time, was not influenced by activation of ACBsh-projecting MCH neurons. Given that depression in humans is generally associated with hyperphagia and body weight gain (Milaneschi et al., 2019), future work is needed to investigate potential interactions between ACB MCH signaling on feeding behavior and depression.

It is well established that there are sex-differences in feeding behavior in rats (Asarian and Geary, 2013), and previous reports indicate that the orexigenic effects of MCH are influenced by sex (Messina et al., 2006; Mogi et al., 2005; Santollo and Eckel, 2008b). For example, in oil- and estradiol-treated ovariectomized female rats, the effect of ICV MCH to increase chow intake was significantly lower in estradiol-treated rats (Messina et al., 2006). Similarly, the capacity of ICV MCH to increase spontaneous meal size is attenuated in estradiol-treated OVX rats (Santollo and Eckel, 2008b). Present results identify a neural site of action for sex- and estradiol-mediated influences on MCH feeding effects. First, using methods that reliably produced hyperphagia in male rats, we observed no effect on feeding following either ACBsh MCH1R stimulation or activation of ACBsh-projecting MCH neurons in intact cycling female rats. Therefore, we hypothesized that the orexigenic effect of ACHsh MCH signaling may be attenuated by estradiol. In bilaterally OVX female rats on a cyclic regimen of either 2 μg EB or oil vehicle, direct injection of MCH in the ACBsh increased feeding only in oil-treated rats, suggesting that EB attenuates the ability of ACBsh MCH signaling to promote chow intake. Our findings thus identify the ACBsh as a likely site of action for the sex-dependent effect of MCH.

The neural mechanisms through which endogenous estrogen influences MCH ACBsh-associated feeding outcomes are unknown. While MCH neurons are in the LHA and ZI in both males and females, prepro-MCH expression has been found in the laterodorsal tegmental nucleus (LDTg) only in female rats (Rondini et al., 2007). Targeting the LDTg MCH neurons in female rats is an area for future exploration. There are also reported sex differences in MCH gene expression, such that prepro-MCH is in the preoptic area in lactating females (Knollema et al., 1992), and more recently, Santollo and Eckel found that both MCH neuronal expression and MCH1R protein expression in the LHA is decreased by estradiol treatment in OVX female rats and in intact cycling female rats (Santollo and Eckel, 2013). Moreover, in cultured hypothalamic neurons, estradiol treatment did not affect MCH or MCH1R mRNA expression, and ERα and MCH-immunoreactive neurons were not co-localized in the LHA/ZI of female rats. Using fluorescence in situ hybridization, we build on these previous findings by revealing similar ACBsh MCH1R mRNA expression in male and female rats. However, MCH1R and ESR1 mRNA expression co-localize in the ACBsh to a greater extent in females than in males. Together, our findings suggest that sex-specific differences in MCH1R and ESR1 mRNA co-localization in the ACBsh may be one of the mechanisms through which estrogen suppresses the orexigenic effect of ACBsh MCH signaling. However, it is possible that MCH1R and ESR1 expression can vary across stages of the estrous cycle. The current stage of estrous at time of tissue harvest was not determined in this experiment and thus should be noted as a limitation of this study.

In conclusion, present data substantiate a role for ACBsh-projecting MCH neurons in promoting feeding in a sex-specific manner and highlight endogenous estrogen signaling as a neurobiological system counteracting the orexigenic response of ACBsh MCH signaling. This orexigenic pathway does not appear to influence incentive motivation for palatable sucrose on a PROG sucrose/chow choice test, general locomotor activity, or anxiety-like behavior. Overall, results contribute to a greater understanding of the role of the MCH system in energy balance regulation and provide novel insight into mechanisms mediating the sexually dimorphic effects of MCH on feeding behavior.

## Author Contributions

SJT, EEN, and SEK designed the research; SJT, KS, RL, CML, and AMC performed experiments; SJT analyzed data; SJT and SEK interpreted results of experiments; SJT and SEK prepared figures; SJT drafted manuscript; SJT, EEN, and SEK edited and revised manuscript; SJT, KS, RL, CML, AMC, EEN, and SEK approved final version of manuscript.

---

**Supplemental Figure 1:**
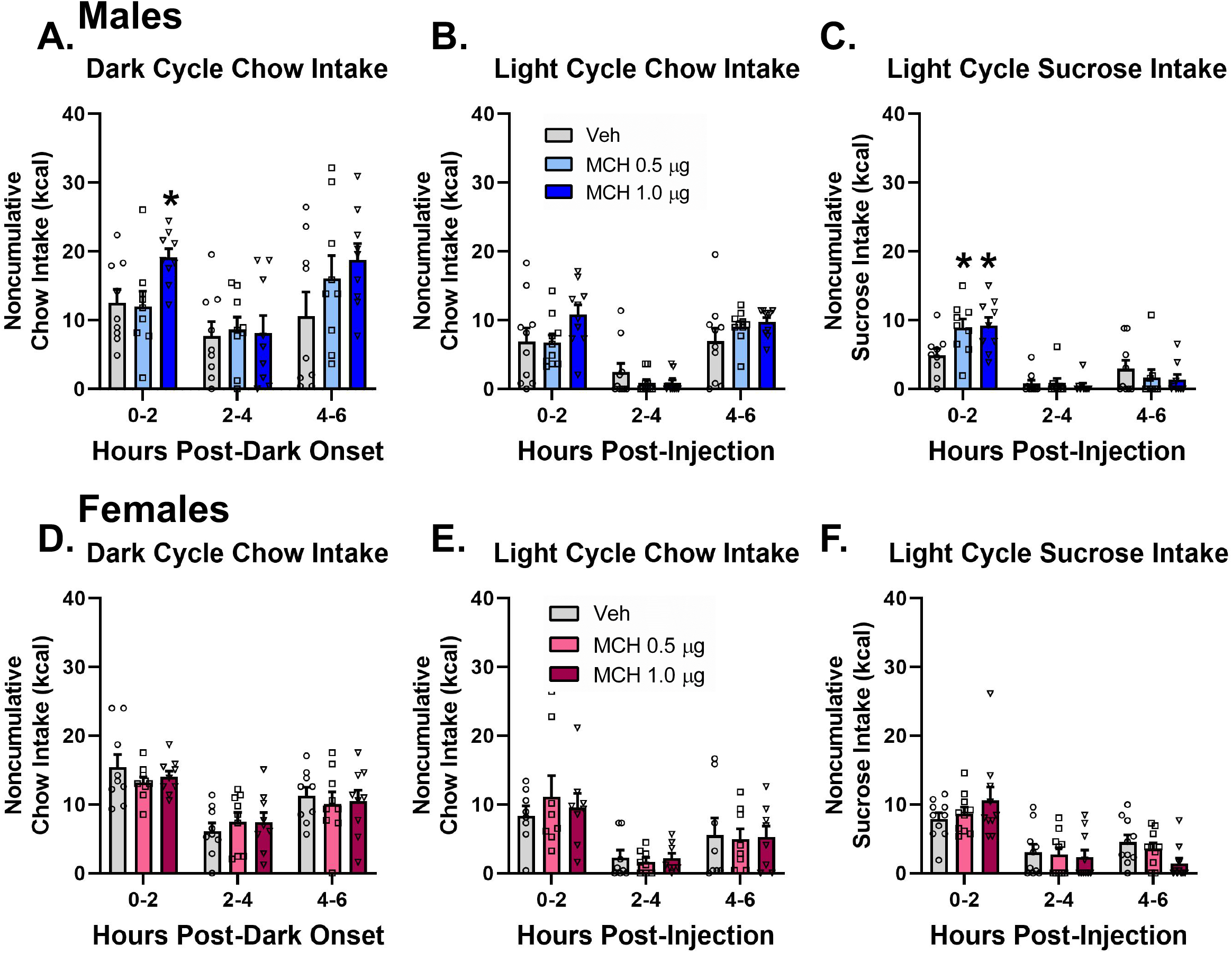
Direct activation of MCH1R in the ACBsh MCH significantly increased noncumulative dark cycle chow intake (a), tended to increase light cycle chow intake (b), and significantly increased light cycle sucrose intake (c) in male rats. ACBsh MCH did not affect food intake during the same noncumulative feeding tests in female rats (d-f). Data are mean ± SEM; *p < 0.05 vs vehicle treatment.

**Supplemental Figure 2:**
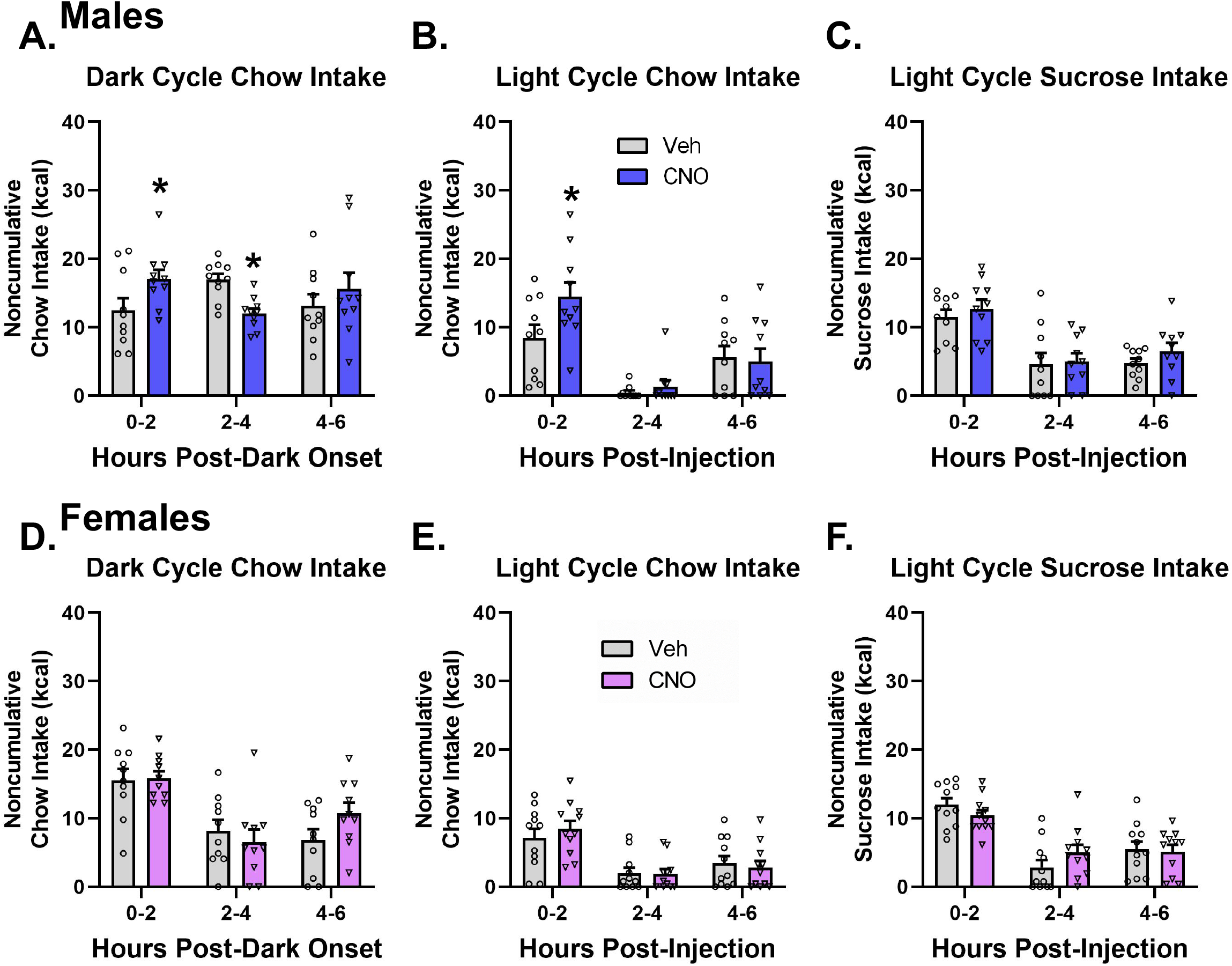
In male rats, activation of ACBsh-projecting MCH neurons following ICV CNO increased noncumulative dark cycle chow intake (a) and light cycle chow intake (b) but did not significantly influence light cycle sucrose intake (c). Activation of ACBsh-projecting MCH neurons had no effect on noncumulative dark cycle chow intake, light cycle chow intake, or light cycle sucrose intake in female rats (d-f). Data are mean ± SEM; *p < 0.05 vs vehicle treatment.

**Supplemental Figure 3:**
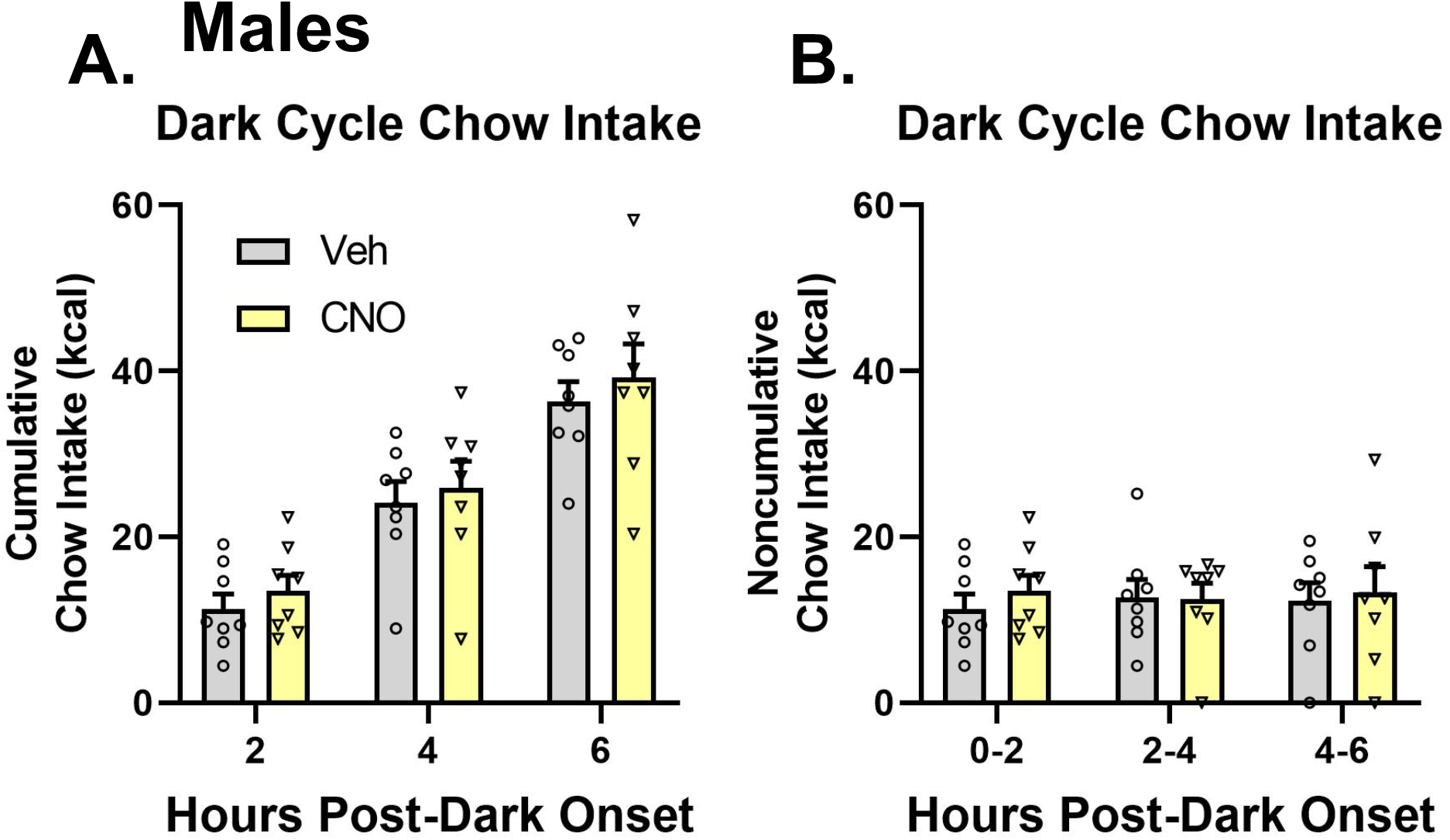
ICV CNO has no effect on either cumulative (a) or noncumulative (b) dark cycle chow intake in male rats lacking DREADDS.

**Supplemental Figure 4:**
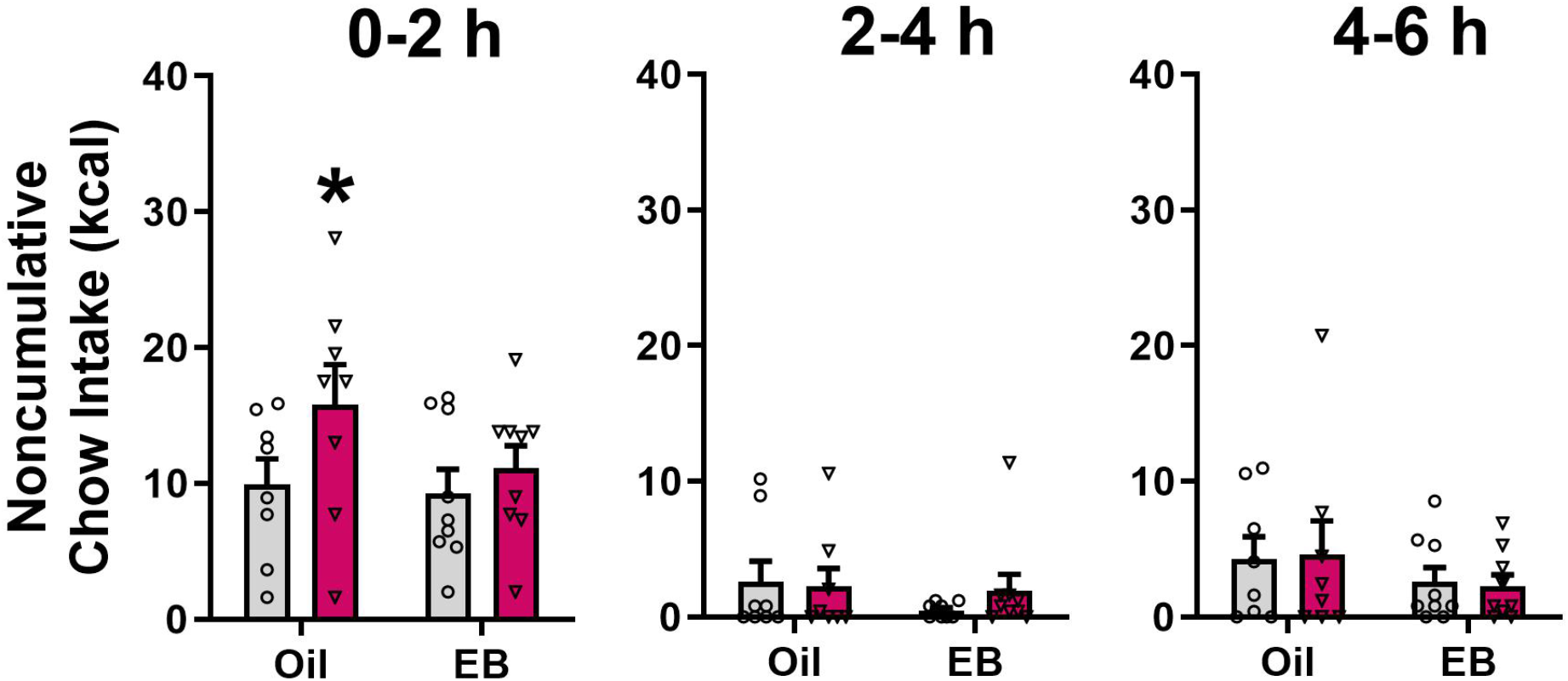
In the oil-treated OVX female rats, 1.0 μg of MCH delivered to the ACBsh significantly increases noncumulative light cycle chow intake during the first 2hr bin; whereas in the EB-treated OVX rats, ACBsh MCH injections did not affect chow intake. Data are mean ± SEM; *p < 0.05 vs vehicle treatment.

